# Regional opportunities for Siberian tundra conservation in the next 1000 years

**DOI:** 10.1101/2021.11.29.470323

**Authors:** Stefan Kruse, Ulrike Herzschuh

**Author notes:** **For correspondence:** (SK).

## Abstract

The biodiversity of tundra areas in northern high latitudes is threatened by invasion of forests under global warming. However, poorly understood nonlinear responses of the treeline ecotone mean the timing and extent of tundra losses are unclear but policymakers need such information to optimize conservation efforts. Our individual based model LAVESI, developed for the Siberian tundra-taiga ecotone, can help improve our understanding. Consequently, we simulated treeline migration trajectories until the end of the millennium, causing a loss of tundra area when advancing north. Our simulations reveal that the treeline follows climate warming with a severe, century-long time lag, which is overcompensated by infilling of stands in the long run even when temperatures cool again. Our simulations reveal that only under ambitious mitigation strategies (RCP 2.6) will ~30% of original tundra areas remain in the north but separated into two disjunct refugia.

## Introduction

Arctic regions have experienced strong warming over recent decades as exemplified in Siberia (***Box et al., 2019***). The warming is expected to continue drastically, based on future changes simulated with prescribed ‘relative concentration pathway’ (RCP) scenarios (***Meinshausen et al., 2011***). Following climate simulations, mean July temperatures in the region could, by the end of the 21st century, be less than +2°*C* warmer with ambitious mitigation efforts (RCP 2.6), about +3.1°*C* (intermediate effort scenario RCP 4.5), or an extreme of +14°*C* warmer (high-emission high-warming RCP 8.5). In consequence, the treeline ecotone can expand and replace tundra (***Hofgaard et al., 2012***; ***MacDonald et al., 2008***). This turnover threatens the existence of various types of sensitive tundra with differing vegetation structure along latitudinal and altitudinal bioclimatic gradients located between the treeline in the south and the Arctic Ocean in the north (***Walker et al., 2005***; ***Pearson et al., 2013***; ***Yu et al., 2009***). The Siberian tundra areas are known for their alpine species biodiversity (***Pauli et al., 2012***; ***Mod and Luoto, 2016***; ***Arctic Climate Impact Assessment, 2004***) and support special (indigenous) types of land use (e.g. reindeer herding, foraging). However, the expansion of forests in response to increasing temperature is not well understood and the uncertain timing of future tundra loss is challenging conservation efforts.

In Siberia, the modelled climate envelope for tundra migrates north by ~ 290 *m yr*^-1^ (***Loarie et al., 2009***). When assuming a direct relationship linking the modern position of the treeline to its main limiting factor of July temperature (***MacDonald et al., 2008***), the treeline is expected to reach its maximum expansion within the first centuries of the millennium. Regionally, where the shoreline of the Arctic Ocean is close to the treeline, the 100-200 *km* wide tundra corridor would be completely covered by forests. Moreover, even regions with wide tundra corridors in Siberia (~ 600 *km*; Taimyr Peninsula and Chukotka) could be forested in the warmest scenarios. However, the potential migration rates are likely constrained by other factors, partly acting on a small scale (***Corlett and Westcott, 2013***). This and the fact that in Siberia the treeline is formed solely by larch species growing on permafrost while in North America this niche is covered by a mix of species (***Mamet et al., 2019**; **Herzschuh, 2020***), make it impossible to transfer knowledge from any other treeline region to inform Siberian tundra responses.

Equilibrium-based estimations are not backed by empirical studies which are showing slower responses and either no treeline advance or rates of only a few metres per year in Siberia (***Scherrer et al., 2020***; ***Kharuk et al., 2013***; ***Wieczorek et al., 2017b***; ***Harsch et al., 2009***). Either the different response of regions can be related to site conditions (e.g. ***Harsch et al., 2009***; ***Mamet et al., 2019***), or it is not well understood how complex processes such as seed dispersal or intra-species competition acting on fine and broad scales constrain the response to warming (***Corlett and Westcott, 2013***; ***Wieczorek et al., 2017b***). Models that do not consider these processes lead to forecasts prone to overestimating treeline advance (compare ***Pearson et al., 2013***; ***Snell and Cowling, 2015***), but it is challenging to consider both small-scale processes and the broad-scale picture.

Models which contain processes to estimate competition between individuals for resources and seed dispersal in a spatially explicit environment and which handle each tree individually are called individual-based spatially explicit models (***DeAngelis and Mooij, 2005***). These models have become increasingly used lately to assist in formulating conservation management strategies (***Zurell et al., 2021***). For the northern Siberian treeline ecotone the vegetation model LAVESI was developed, which incorporates a full tree (*Larix*) life-cycle beginning with seed production and dispersal, germination, and establishment of seedlings which then grow based on abiotic (temperature, precipitation, active layer depth) and biotic (competition) conditions (***Kruse et al., 2019a**, **2016***). Although soil nutrients and mycorrhiza are not considered explicitly in the current version, it was shown that it can capture the nonlinear response of the treeline populations well at a regional level on the Taimyr Peninsula (***Kruse et al., 2019a**, **2016**; **Wieczorek et al., 2017b***).

To cope with the complex forest-tundra ecotone responses, which are not in equilibrium with climate (***Hofgaard et al., 2012***), we updated and made use of our individual-based spatially explicit model LAVESI. With our simulation study, we explore treeline migration and tundra area dynamics to provide important information as to which tundra areas can survive future warming and where preventive measures for protection would be necessary to conserve the unique Siberian tundra. This study is guided by two questions: **“What are the treeline trajectories in Siberia for different climate mitigation scenarios (RCP)?”** and **“Under which climate scenario can tundra retreat to refugia and recolonize areas after temperatures cool again, and where?”**

## Results and Discussion

### Treeline migration trajectories under future climate changes

The simulations revealed that treeline advance follows climate warming exponentially reaching 30 *km decade*^-1^ in the 21st century with faster rates the warmer the climate scenario (***Table 1***). When relating and tracking the position of the treeline to the climate (= climate-analogue) we find a severe migration lag of hundreds of years until simulations reach the same position (***Figure 1***). This behaviour is longer than expected from the properties of treeline-forming larch species with generation times of decades and weak long-distance dispersal rates (e.g. ***Mamet et al., 2019***; ***Kruse et al., 2019a***; ***Scherrer et al., 2020***; ***Clark, 1998***). Other processes, not yet explicitly captured by the model, may constrain the migration process, such as the strong greening of the Arctic (compare ***Pearson et al., 2013***; ***Berner et al., 2020***), the warming and abrupt thaw of permafrost (e.g. ***Stuenzi et al., 2021***; ***Biskaborn et al., 2019***), soil nutrients (e.g. ***Sullivan et al., 2015***), and animal activity (e.g. ***Wielgolaski et al., 2017***). Regardless, our simulated rates over the past few decades are generally in line with the few existing modern observations showing only a slow treeline advance (***Mamet et al., 2019***; ***Hansson et al., 2021***; ***Rees et al., 2020***; ***Shevtsova et al., 2020***; ***Guo and Rees, 2021***). Rates are especially high in Chukotka, however, in all forcing scenarios (***Table 1***): an observation that can be attributed to forest patches ahead of the current treeline, which serve as nuclei for tundra inflling (shown by e.g. ***Snell and Cowling, 2015***; ***Kharuk et al., 2013***; ***Kruse et al., 2019a***; ***Harsch and Bader, 2011***).

After the initial migration lag we can see from our millennium-long simulations that the treeline reaches more advanced positions than the climate analogue (***Figure 1**, **Table 1**, **Appendix 1 Figure 1**, **Appendix 1 Figure 2**, **Appendix 2 Figure 1**, **Appendix 2 Figure 2***). This strong overshooting was unexpected especially in scenarios with cooling temperatures back to the 20^*th*^ century level. However, we designed our model to be pattern-oriented and is based on observations that, once established, trees can endure periods of unfavourable environments (compare ***Kruse et al., 2016***). This leads to the survival of trees ahead of the treeline acting as nuclei for tundra colonization when conditions improve (***Väliranta et al., 2011***; ***Snell and Cowling, 2015***). This is backed by observations of slower than expected retreat of elevational treelines because trees can persist until dramatic events cause the death of all well-established trees (***Scherrer et al., 2020***).

**Figure 1.**
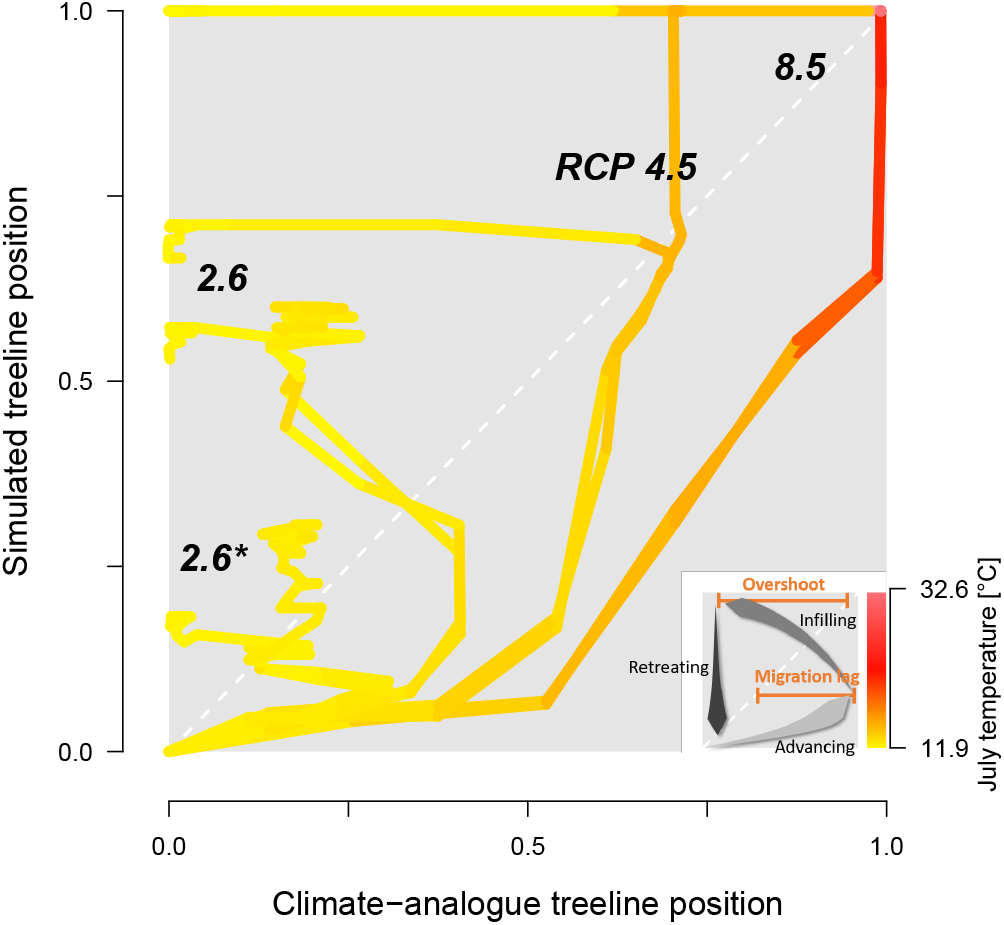
The trajectory of the simulated treeline position versus its current position for the Taimyr Peninsula shows a migration lag of the treeline during the first centuries (each line segment represents 25 years) until the simulated treeline is limited by climate. Forests expand their area further and infilling proceeds when climate conditions cool and even overshoot in the long run with cooling back to *20^th^* century temperatures.

### Tundra area dynamics caused by forest expansion

The area loss over time roughly follows a negative exponential function, which is steeper the warmer the climate scenarios is (***Figure 2**, **Appendix 2 Figure 1***). The quickest loss, already by - 2100 *CE*, is seen in the intermediate warming scenario RCP 4.5 (+3 °*C* July temperatures) and the extreme RCP 8.5 (July temperatures +14 °*C*). For these scenarios, tundra retreats to its minimum cover of 5.7% by the middle of the millennium (Fig. 2b). Even under mitigation scenario RCP 2.6 the tundra area is reduced to 32.7% or in the very ambitious scenario 2.6* to only 54.9%. Other simulation studies show 50% loss during the 21st century (***Callaghan et al., 2005***; ***Kaplan et al., 2003***; ***Pearson et al., 2013***) but not the dynamics of the long run until the end of 3000 *CE*, as shown by our study.

**Figure 2.**
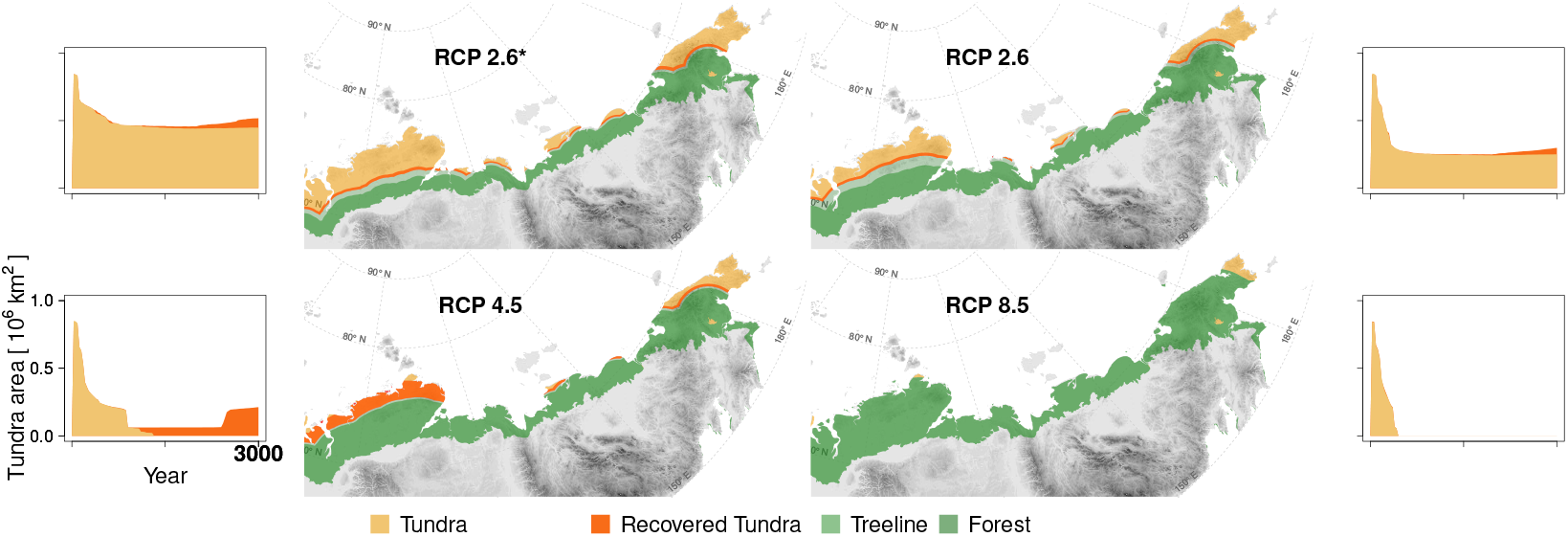
Forest position and tundra area at year 3000 *CE* for different climate mitigation scenarios and under potential cooling back to *20^th^*century temperatures after peak temperatures have been reached. The area of tundra changes over time and can only partly recover when temperatures cool and forests recede (plots next of the maps show years 2000-3000 *CE*). Only tundra areas above the treeline (***Walker et al., 2005***) are considered;Map projection: Albers Equal Area. **Figure 2–video 1.** The development of the forest position and tundra area can be seen in a video supplement presenting the state in 10-year steps.

Cooling back to the 20^*th*^ century levels led to a slight tundra re-expansion of a few percent of the area by the end of the simulations. While re-expansion is small for RCP 2.6* and 2.6, and largest for RCP 4.5 on the Taimyr Peninsula where forests retreat from areas of ~ 200 000 *km*^2^ in the final 200 years of the simulation, under RCP 8.5 the effect of the cooling led to no tundra releases. In consequence, only if humans act ambitiously in limiting global warming to well below +2°*C* (RCP 2.6, 2.6*), and temperatures actually cool after the maximum, do simulations show that ~10-30% of the tundra area could persist in the north and can re-expand after the mid-millennium maximum treeline extent. However, due to the overshooting effect, less than 10% of the area remains unforested. All in all, the expansion is in accordance with other studies and will likely put especially cold-climate tundra types at risk of extinction (e.g. ***Pearson et al., 2013***; ***Walker et al., 2005***).

### Concluding opportunities for tundra conservation

Following our simulation results, tundra is reduced to ~5.7% of its current area at maximum forest expansion under the most likely scenario RCP 4.5 after an initial migration lag. Only ambitious strategies to mitigate climate warming (for example RCP 2.6) could prevent such an extensive loss and retain ~32.7% of the current tundra. In both cases, the northern refugia are located in Chukotka and the Taimyr Peninsula, currently covered by High Arctic and Arctic tundra types which have relatively high biodiversity (***Arctic Climate Impact Assessment, 2004***). A WWF report (***Krever et al., 2009***) finds that the current total cover of protected northern areas is insufficient to achieve the requisite 30% protection necessary for biodiversity conservation (***Secretariat of the United Nations Convention on Biological Diversity, 2021***). Especially cold-climate tundra types on Siberian islands and in Chukotka are under-protected (e.g. ***Pearson et al., 2013***; ***Walker et al., 2005***) and thus further protected areas need to be established.

Although the large refugia could enable tundra species survival and later recolonization (analo-gon forests, ***Harsch et al., 2009***; ***Clark, 1998***; ***Stralberg et al., 2020***), the diversity in these ~ 2500 *km* distant fragments is threatened by the fundamental disadvantages of a small population (inbreeding/genetic drift, ***Ohsawa and Ide, 2008***; ***Charlesworth and Charlesworth, 1987***; ***Mona et al., 2014***; ***May et al., 2013***). A network of connected systems would be necessary with protection of potential refugia (e.g. ***Morelli et al., 2020***) distributed across the ~ 4000 *km* wide tundra area. Due to its individual-based nature, LAVESI can be a tool to assess population genetics and aid in the optimal placement of migration corridors and areas to protect (cf. ***Zurell et al., 2021***).

## Methods

### Model description and improvements

The model LAVESI is an individual based and spatially-explicit model that was developed to study the treeline ecotone growing on permafrost soils on the Taimyr Peninsula (***Kruse et al., 2016***; ***Wieczorek et al., 2017b***; ***Epp et al., 2018***) and since then it has been improved and modified for different purposes including for migration-rate studies (***Kruse et al., 2019a***). To achieve the most realistic model, each life-history stage of the larch individuals is handled explicitly, and the model’s processes adapted to observed patterns of surveyed tree stands for the dominant larch tree species, *Larix gmelinii* and *Larix sibirica*. The source code was updated to version 1.2 on the crutransects branch to be sufficiently light in terms of memory allocation and include landscape (developed in parallel for Chukotka mixed alpine-latitudinal treeline simulations in ***Shevtsova et al., 2021***; ***Kruse et al., 2021***). The model is freely available on Github https://github.com/StefanKruse/LAVESI. The source code of the version used in this manuscript will be additionally permanently stored on Zenodo on final publication.

### Model performance

During simulation runs, each Environment grid cell of *20×20 cm* needs 10 *bytes*, an element of the Tree-structure 64 *bytes* and the Seed-structure 32 *bytes*. In total, for very dense scenarios it needs for each square km ~ 11 *GB RAM*. The parallelization was updated and instead of Standard Template Library (STL) list containers for tree and seed elements a vector structure was implemented to allow better support for OpenMP (https://www.openmp.org) parallelization for scaling on a High-Performance Cluster and the realization of large-scale transect simulations as needed for this study. The computation speed depends on the number of trees and reaches ~ 3.5 *s yr*^-1^ *km*^-2^ for very dense forests on 8 cores that can be reduced when using more cores ***Figure 3***. However, due to overheads, scaling is not 1:1. Of the internal functions of LAVESI the mortality is most computationally intensive, as each seed and tree need to be handled individually.

**Figure 3.**
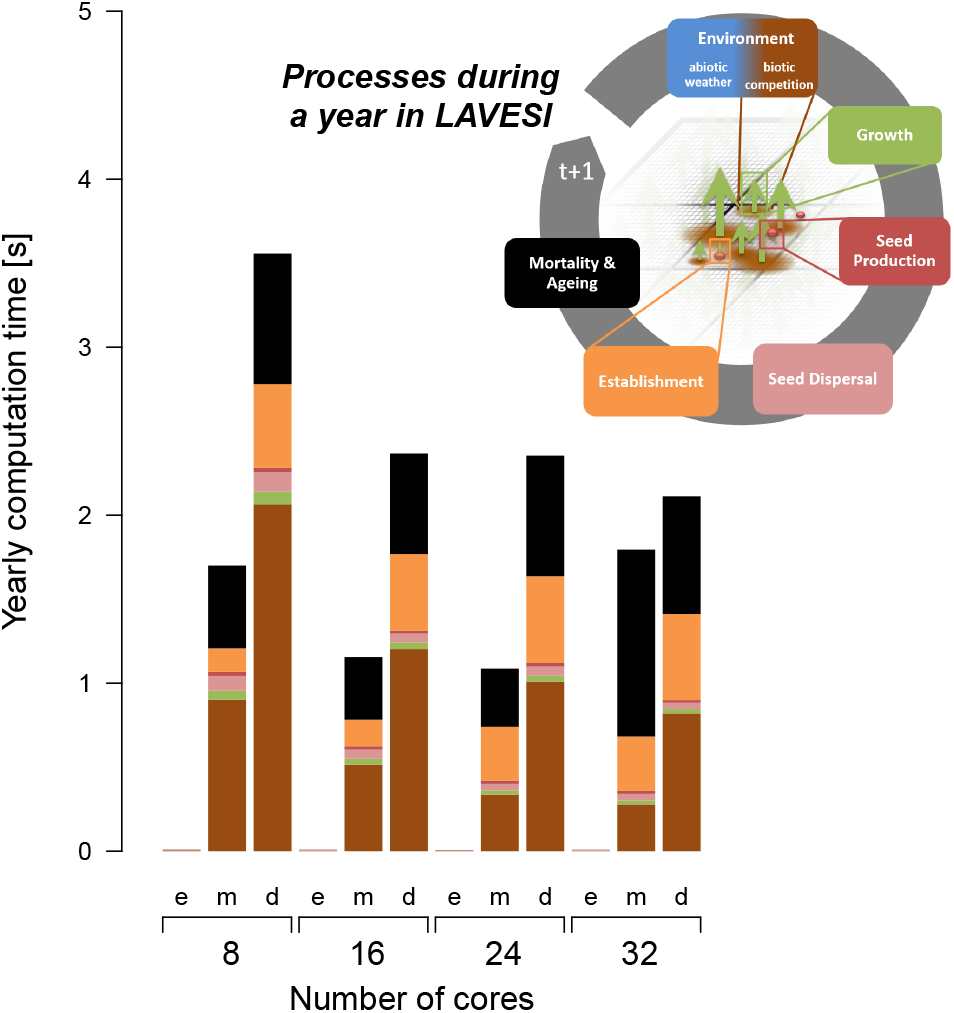
The computation speed increases with the number of available threads and reaches a plateau caused by overheads. The processes of LAVESI scale differently and the process Environment is computationally the heaviest and thus relatively faster with more cores while processes such as Mortality show no clear benefit of more threads. Data were aggregated for 100 years at three different population dynamics stages: e=empty, m=mature old growth, d=densification.

### Transect setup

At four representative locations of the Siberian treeline within 100 – 170 °*E*, we placed linear transects, starting at a field site, and allowed open tree stands to establish upto the shoreline (***Figure 4***). From west to east: (1) Taimyr Peninsula, 573 *km* long, site 13-TY-09-VI, 72.15067° *N* 102.09771° *E* (***Wieczorek et al., 2017b**,c*), (2) Buor Khaya Peninsula, 137 *km* long, 14-OM-20-V4: 70.52671° *N* 132.91426° *E* (***Liu et al., 2020***), (3) Kolyma River Basin, 146 *km* long, 12-KO-02/I: 68.38916° *N* 161.46617° *E* (***Wieczorek et al., 2017a***), (4) Chukotka, 626 *km* long, EN18022: 67.40102° *N* 168.34801° *E* (***Kruse and Stoof-Leichsenring, 2016***; ***Kruse et al., 2019b***). While the first three locations in the west cover predominantly flat terrain, the easternmost location in Chukotka has mountainous stretches.

**Figure 4.**
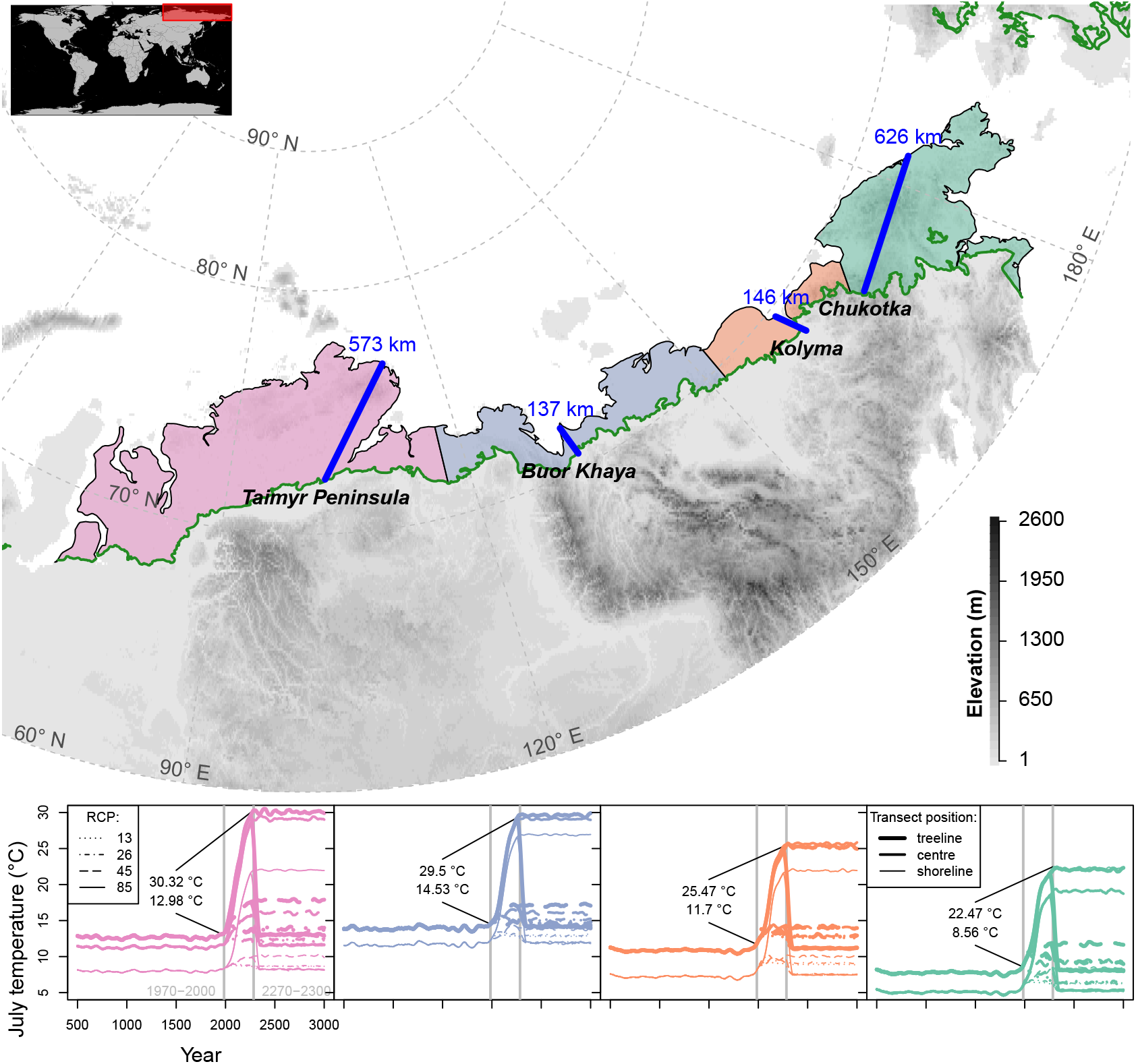
Transects (blue lines) were placed starting in the treeline at field sites and estending to the shoreline of the Arctic Ocean (map); only tundra areas above the treeline (***Walker et al., 2005***) are considered. The land area was grouped into four regions equidistantly separated between the transects (lower plot) and plots show climate forcing for 501-3000 *CE* based on RCP scenarios in these regions at the extremes and middle part of the transects; numbers give mean values for modern and future strongest warming under RCP 8.5. Map projection: Albers Equal Area.

### Climate forcing

Temperature and precipitation time series were constructed using the following steps:

1. Extract climate data in 10 *km* steps along transects beginning at the position of the field sites for each of the four focus regions from the grid cells of CRU TS 4.03 by distance weighted interpolation (***Harris et al., 2020***).
2. Establish a yearly *5*^18^O and ice layer thickness series for 501 to 1900 from the yearly resolution ice-core data from Severnaya Zemlya (north of the Taimyr Peninsula, available for years 934 to 1998 ***Opel et al., 2013***), the only closest annual resolution archive allowing both temperature and precipitation reconstructions independent of plant-based proxies (pollen or tree rings). This was achieved by sampling 25-year long blocks of yearly resolution data from the pre-industrial (< 1800 *CE*) and adding this variability to the AN86/87 core which dates back to 4703 *BCE* (***Arkhipov et al., 2008***).
3. Adjust this locally to the 10 *km* steps climate data series per transect by building linear models of yearly temperature and precipitation for the overlapping years 1901-1998 and use this to estimate mean values for the period 501 to 1900.
4. Sample 15-year long blocks from CRU and adjust the mean temperature by the difference of the sampled block from the estimated mean values to achieve monthly data series.
5. Prolong this series by coupled climate model long runs until 2300 *CE* with model output with MPI-ESM-LR prepared for CMIP5 for scenarios RCP 2.6, RCP 4.5, and RCP 8.5 and additionally a scenario that warms only half the values of RCP 2.6, which is here called RCP 2.6*.
6. Extend these trends for 2100-2300 until 2500 *CE* and repeat in the following either the climate data of 1901-1978 in a loop which in this study is called *20^th^* century cooling or copy the period 2300–2500 until 3000 *CE*.

July temperatures (***Figure 4***), which have the strongest impact in the model, increase in these scenarios by a median of +1.2 °*C* (RCP 2.6*), +1.8 °*C* (2.6), +2.1 °*C* (4.5) and up to +5 °*C* (8.5) compared to 1970-2000 until 2100, which partly cools until 2300 to +0.6 °*C* (RCP 2.6*), +0.5 °*C* (2.6) and reaches higher levels for the two warmest scenarios to +3.1 °*C* (4.5) and up to +14.4 °*C* (8.5).

Wind speed and direction data were obtained at 6-hourly resolution for 1979-2018 from EraIn-terim5 (data set description in ***Dee et al., 2011***) for each of the transects and supplied with the model. In a prior version, these data were only available for the Taimyr Peninsula and only for years until 2012.

### Tree growth on transects

We implemented sensing of the environment for each individual tree via its y-position along the transect which is made up of 10 *km* spaced climate data. The maximum possible growth at a certain position (see initial publication ***Kruse et al., 2016***) is calculated by interpolating the possible growth under the climate using the two closest climate variables.

The larch species *Larix gmelinii* (Rupr.) Rupr. dominates the areas to the west of the Verkhoyansk Mountain Range ~90-130° E and its sister species *L. cajanderi* Mill. grows in north-east Siberia ***Figure4*** (***Abaimov,2010***). Hence, the already implemented species *L.gmelinii* was set to be present in the simulations and simulations tuned to fit observations. To initiate growth 100 000 *seeds* were introduced at the cold end with a distance based on a negative exponential kernel for the first 50 *years*. Similarly, extinction on a transect was prevented by permanently introducing 1000 *seeds* per 200 *km* length. Further seed introduction from the non-explicitly simulated hinterland was estimated on a 500 *m* long stretch at the cold site linking the production and release height to climate.

### Model tuning and validation

Simulation runs were forced with the prepared climate data for the transects for validation of the shape of the treeline as simulated by the model. The results of year 2000 were compared to field data that was positioned along the temperature gradient on the simulated transects by using the respective CRUTS temperature data for each grid cell of each plot (example ***Figure 4***). The field data collection included disturbed sites (e.g. in Chukotka) leading to very high stem counts, which are the number of trees exceeding 1.3 *m* in height per ha. The model could be tuned by introducing a local adjustment based on elevation lapse rates of the relevant climate values used internally, which is necessary as the climate data may lead to over/underestimations locally in a model that does not consider local topography: an issue that is especially relevant in the mountainous region of Chukotka. We made a moderate fit with the following amendments

- Taimyr Peninsula: *T_Jan_* = −1.17 °*C*, *T_JuI_* = −0.84 °*C* and *P_Year_* = −5.4 *mm*
- Buor Khaya Peninsula: *T_Jan_* = −0.23 °*C*, *T_Jul_* = −2.48 °*C* and *P_Year_* = −28.9 *mm*
- Kolyma River Basin: *T_Jan_* = +3.14 °*C*, *T_Jul_* = +1.26 °*C* and *P_Year_* = +85.7 *mm*
- Chukotka: *T_Jan_* = +4.46 °*C*, *T_Jul_* = +4.30°*C* and *P_Year_* = +8.2 *mm*

From the individuals present along the simulated transects, the northernmost positions of the treeline are calculated. For the four focus regions these are +66, +88, +98, and +16 *km* beyond the treeline start location at year 2000 (***Appendix 1 Figure 2***), which are partly ahead of the observed positions based on visual inspection of satellite data ~ +30, +30, +80, and +15 *km* beyond the field location selected as the starting point.

### Simulation setup and data processing

We ran three simulation repeats for each transect with the climate series prepared for this study as initial tests showed robust simulation results. To consider the available area from the start to the shoreline, we used 800 *km*-long and 300 *km*-long transects and 20 *m* wide wrapping for the east and west boundary of each simulated stretch, thereby representing a slice of a continuous treeline.

The positions of three key variables are extracted in 10-year steps: single-tree stands, treeline stands, and forestline stands, which are defined as the northernmost position of stands with > 1 *stem* (tree > 1.3 *m* tall) per ha, the northernmost position of a forest cover not falling below 1 *stem ha*^-1^, and the northernmost position of a forest cover not falling below 100 *stems ha*^-1^ (see Fig. 2 in ***Kruse et al., 2019a***, for a graphical representation), respectively. The determined treeline at year 2000 is used as a baseline for expansion and subtracted from the following years’ values. The resulting values for each of the three key variables are used in 10-year steps to interpolate with a weighted average the expansion starting at the current treeline (e.g. ***Walker et al., 2005***).

We used R version 3.6.1. (***R Core Team, 2019***) for data handling, statistical analyses and graphical representation. The 30 arc sec WorldClim 1.4 data (***Hijmans et al., 2005***) along the transects and 2.5 minutes of a degree elevation for mapping was downloaded through the geospatial package raster version 3.0-12. (***Hijmans, 2020***). Additional geospatial packages sf version 1.4-2 and rgdal version 1.4-7, (***Pebesma, 2018***; ***Bivand et al., 2019***) were used to reproject the treeline shape to Albers Equal Area projection and calculate weighted average buffers at each 10 *km*-step along the treeline and merge the buffer polygons for each of the advancement values for the three key variables (single trees, treeline, forestline). The treeline constructed by the circumarctic vegetation map consortium (CAVM, ***Walker et al., 2005***) is provided in segments that were simplified in QGIS version 3.10 (***QGIS Development Team, 2020***). For tundra area assessment, the land area polygons of Russia accessed via R package maps version 3.3.0 (***code by Richard A. Becker et al., 2018***) were cleaned and the islands in the Arctic Ocean excluded. Linear models for temperature and precipitation reconstruction and climate lapse rates were established with functions from the R base package (***R Core Team, 2019***).

## Acknowledgments

This studyacknowledges support by the ERC consolidator grant(no. 772852) led by Ulrike Herzschuh. We thank the logistics team of AWI and the North Easter Federal University in Yakutsk (NEFU) for expedition support and all colleagues that helped during the fieldwork. We like to thank Sven Will-ner for assisting in programming and OpenMP improvements and Cathy Jenks for proofreading and improving the manuscript.

## Appendix 1 Migration rates

***Appendix 1 Table 1*** provides a summary of simulated treeline advance and the corresponding position at the same year of the climate analogue. ***Appendix 1 Figure 1*** shows the position of the climate-analogue treeline position at the four transects and ***Appendix 1 Figure 2*** gives an overview of the simulated maximum positions of the single-tree stands, the treeline and the forestline at the same transects.

**Appendix 1 Figure 1.**
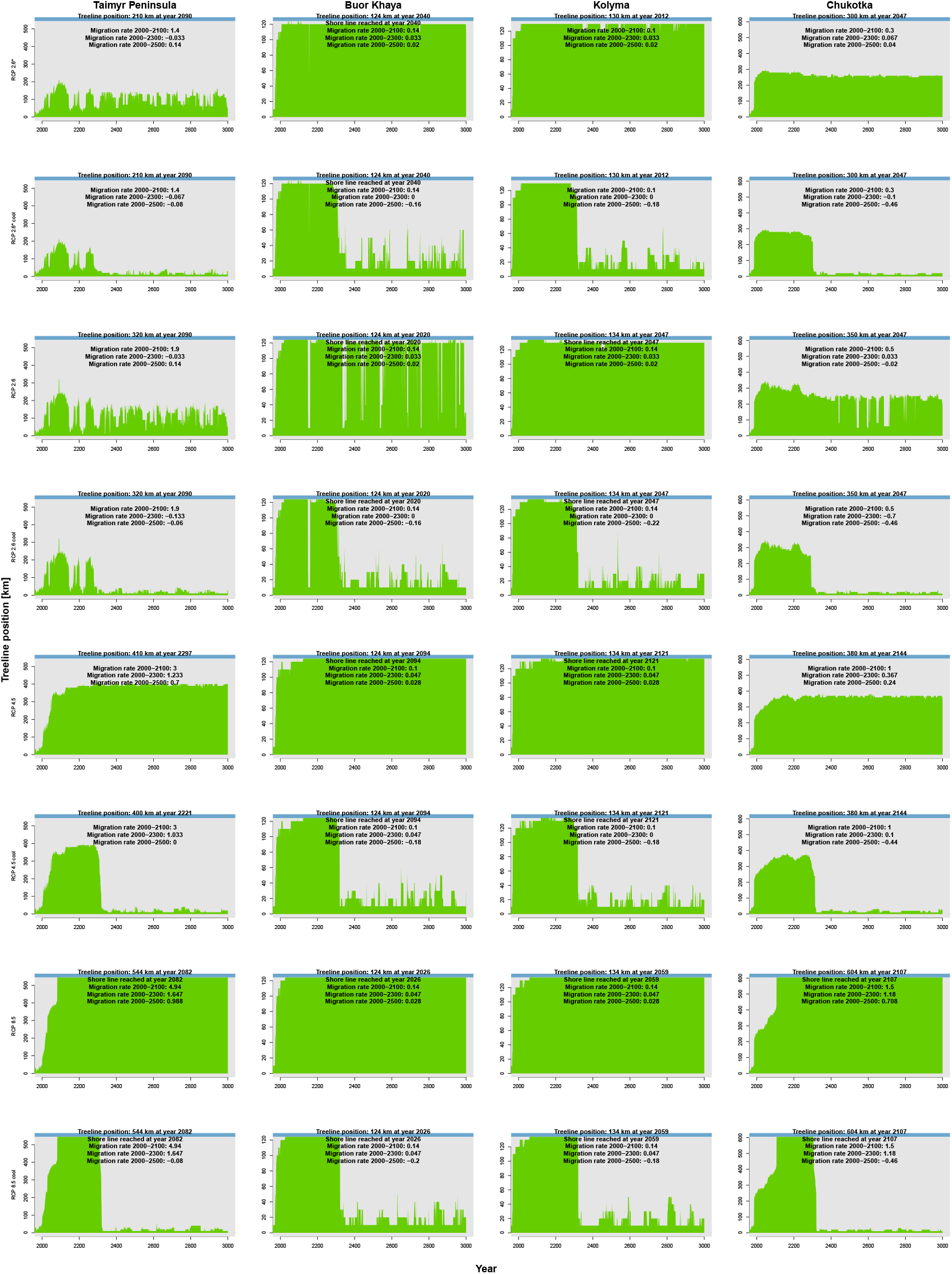
Forest expansion dynamics along the four transects (columns) extrapolated based on July temperatures (climate-analogue) for climate data based on RCP scenarios (rows). A general rapid expansion of the forestline (continuous cover from south > 1 *tree ha*^-1^, green) can be seen. Blue colour at the top represents the shoreline.

**Appendix 1 Figure 2.**
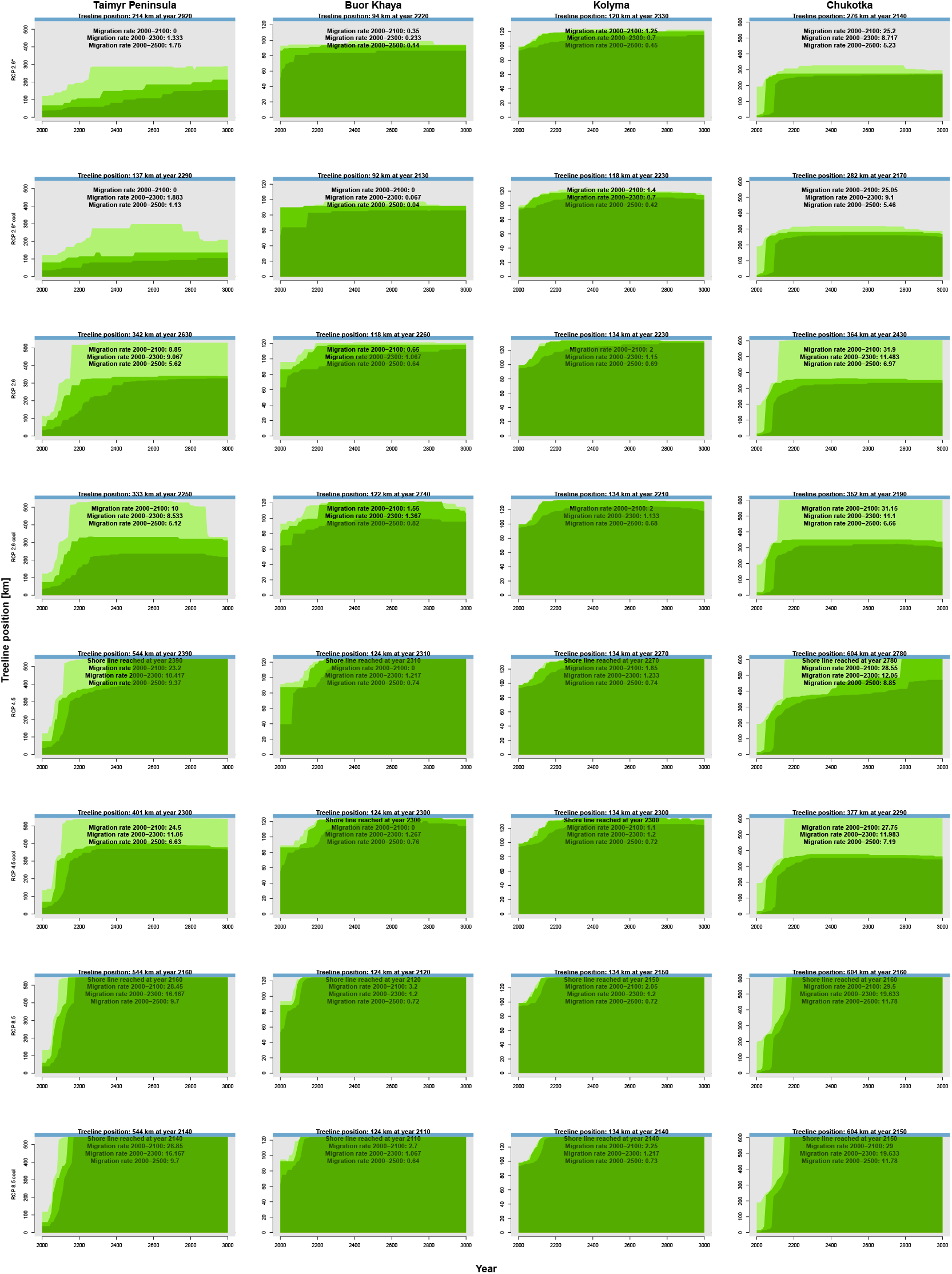
Forest expansion dynamics along the four transects (columns) forced with climate data based on RCP scenarios (rows). A general quick expansion of single tree stands (maximum position of > 1 *tree ha*^-1^, light green) and the forestline (continuous cover from south > 1 *tree ha*^-1^, green) followed by the treeline (continuous cover from south > 100 *tree ha*^-1^, dark green) is seen. Blue colour at the top represents the shoreline.

**Appendix 1 Table 1.**
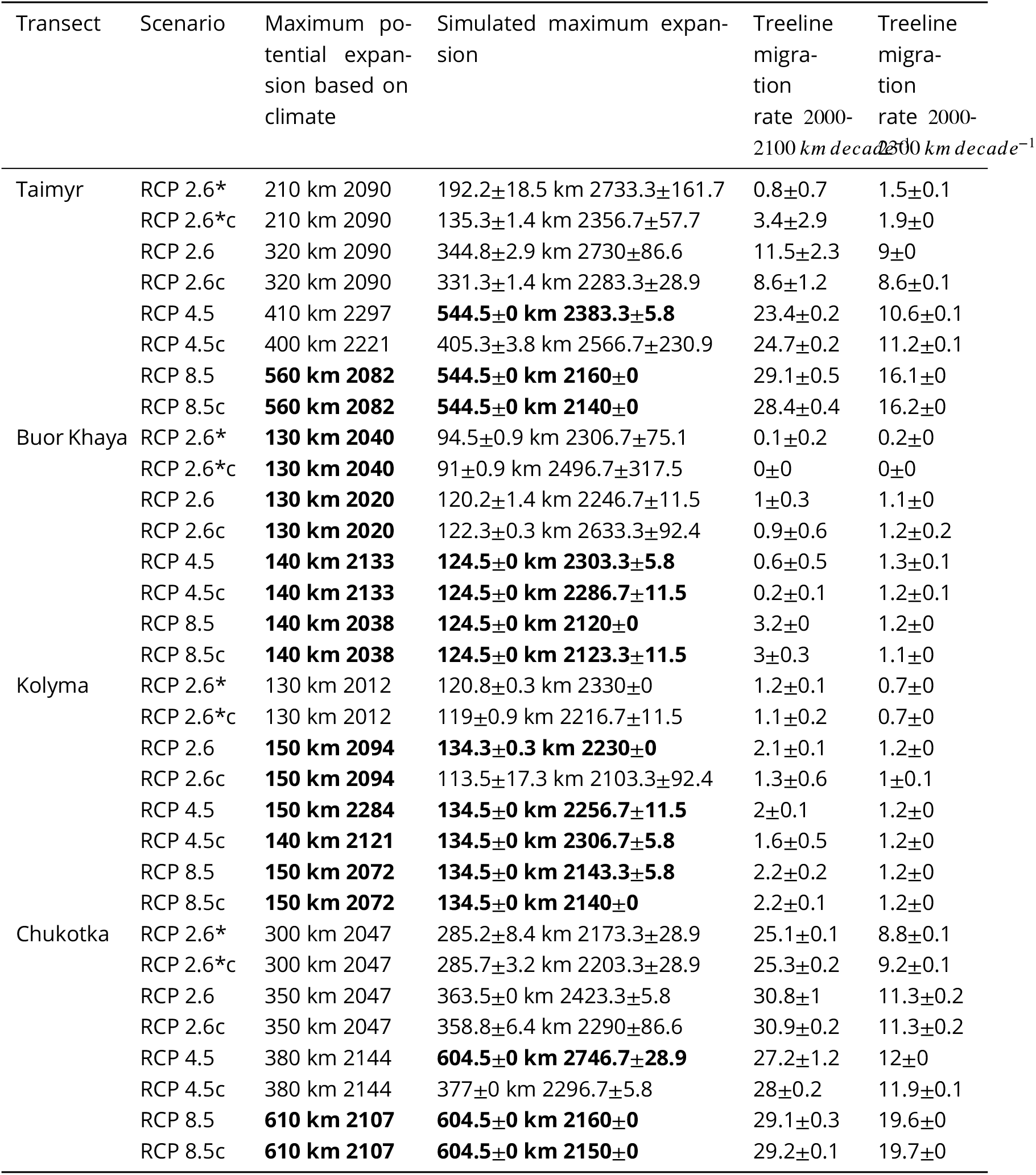
Summary of simulated vs. climate envelope-based treeline advance. Climate envelope solely based on July temperatures (following ***MacDonald etal., 2008***). Values in bold reach the shoreline, but note that because of technical reasons the step sizes of the climate are in 10 *km* steps.

**Appendix 2 Figure 1.**
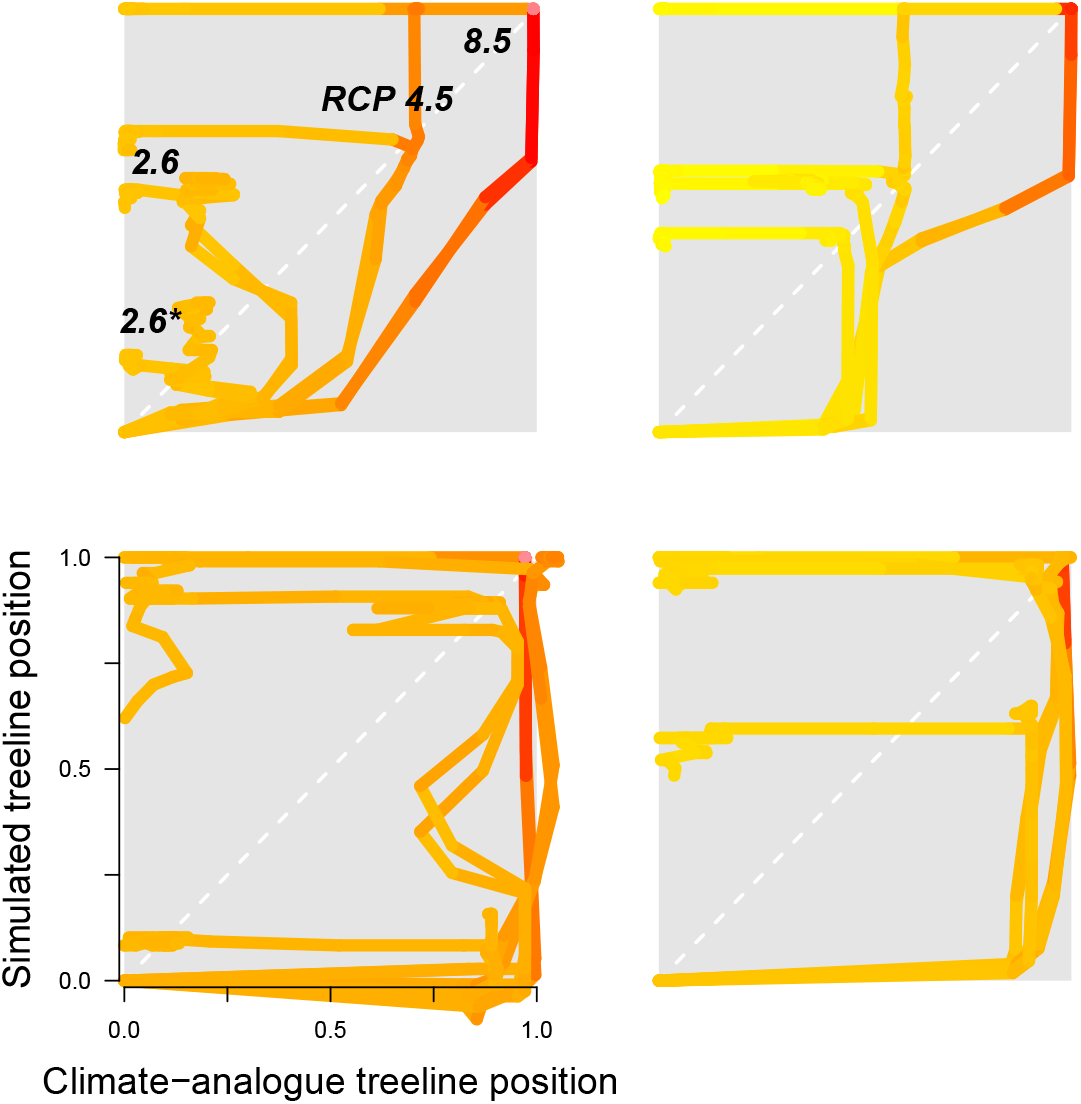
Trajectories for all four regions. Numbers are the first year when the simulated treeline position is equal to or farther north than the modern climate-analogue position. Colour of line segments ranges from yellow for year 2000 to blue in 3000 *CE*. Further description in caption of ***Figure 1*** in main text.

## Appendix 2 Treeline migration trajectories

***Appendix 2 Figure 2*** shows the treeline trajectories of the position based on simulations and the climate-analogue position at the four transects and the time when the simulated positions reach the first time the same position of the climate-analogue. ***Appendix 2 Figure 1*** includes additionally the information of the current temperature in 25-year segments.

**Appendix 2 Figure 2.**
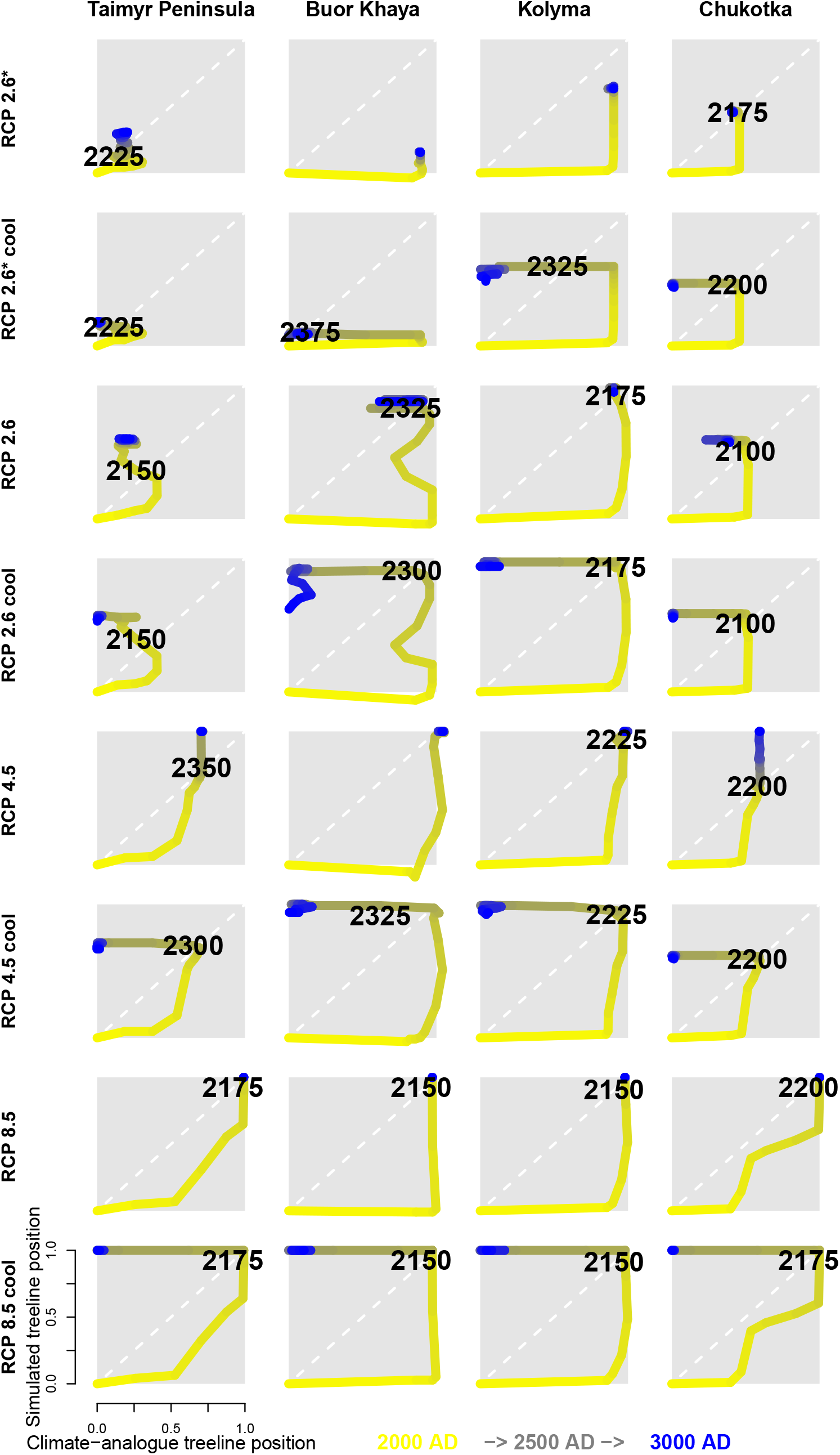
Trajectories for all four regions clockwise: Taimyr Peninsula, Chukotka, Kolyma, and Buor Khaya Peninsula. Colour of line segments ranges from yellow for year 2000 to red in 3000 or blue in year 3000 under the cooling scenario. Further description in caption of ***Figure2*** in main text.

## Appendix 3 I Tundra area dynamics

***Appendix 3 Figure 1*** gives an overview of the tundra area development in 100-year time steps for each of the eighth climate scenarios.

**Appendix 3 Figure 1.**
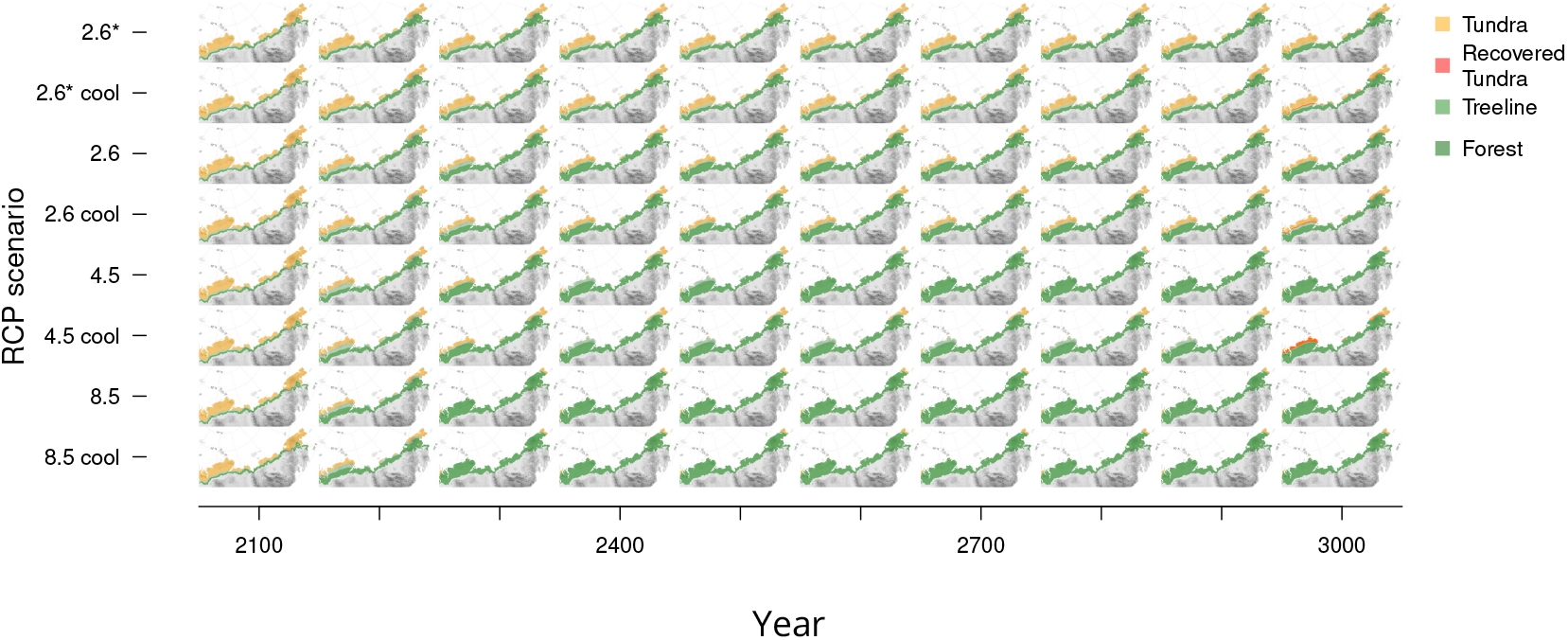
Forest expansion for 100-year time steps and all scenarios. Modern tundra areas will become covered by at least open larch tundra forests under different climate scenarios and nearly reach the shoreline in warmest conditions.

## References

Abaimov AP. Geographical Distribution and Genetics of Siberian Larch Species. In: Osawa A, Zyryanova OA, Matsuura Y, Kajimoto T, Wein RW, editors. Permafrost Ecosystems - Siberian Larch Forests, vol. 209 of Ecological Studies Dordrecht: Springer Netherlands; 2010.p. 41–58. doi: 10.1007/978-1-4020-9693-8.

Arctic Climate Impact Assessment. Impacts of a Warming Arctic-Arctic Climate Impact Assessment. Cambridge University Press, Cambridge, United Kingdom and New York, NY, USA; 2004.

Arkhipov SM, Kotlyakov V, Punning YMK, Zogorodnov V, Nikolayev VI, Zagorodnov VS, Macheret YY, Vaikmaye R, Barkov NI, Korsun SA, Korotkevich V, Morev VA, Evseyev AV, Vostokova TA, Andreev AA, Klementyev OL, Korotkevitch YS, Stiévenard M, Sinkevich SA, Samoylov OY, et al., Deep drilling of glaciers: Russian projects in the Arctic (1975-1995). PANGAEA; 2008. doi: 10.1594/PANGAEA.707363.

Berner LT, Massey R, Jantz P, Forbes BC, Macias-Fauria M, Myers-Smith I, Kumpula T, Gauthier G, Andreu-Hayles L, Gaglioti BV, Burns P, Zetterberg P, D’Arrigo R, Goetz SJ. Summer warming explains widespread but not uniform greening in the Arctic tundra biome. Nature Communications. 2020 dec; 11(1):4621. doi: 10.1038/s41467-020-18479-5.

Biskaborn BK, Smith SL, Noetzli J, Matthes H, Vieira G, Streletskiy DA, Schoeneich P, Romanovsky VE, Lewkowicz AG, Abramov A, Allard M, Boike J, Cable WL, Christiansen HH, Delaloye R, Diekmann B, Drozdov D, Etzelmüller B, Grosse G, Guglielmin M, >et al. Permafrost is warming at a global scale. Nature Communications. 2019; 10(1):1–11. doi: 10.1038/s41467-018-08240-4.

Bivand R, Keitt T, Rowlingson B. rgdal: Bindings for the ‘Geospatial’ Data Abstraction Library; 2019, https://cran.r-project.org/package=rgdal.

Box JE, Colgan WT, Christensen TR, Schmidt NM, Lund M, Parmentier FjW, Brown R, Bhatt US, Euskirchen ES, Romanovsky VE, Walsh JE, Overland JE, Wang M, Corell RW, Meier WN, Wouters B, Mernild S, Mård J, Pawlak J, Olsen MS. Key indicators of Arctic climate change: 1971-2017. Environmental Research Letters. 2019 apr; 14(4):045010. https://iopscience.iop.org/article/10.1088/1748-9326/aafc1b, doi: 10.1088/1748-9326/aafc1b.

Callaghan TV, Björn LO, Chernov YI, Chapin III FS, Christensen T, Huntley B, Ims R, Johansson M, Jolly D, Matveyeva N, Panikov N, Oechel W, Shaver G. Arctic tundra and polar desert ecosystems. Arctic Climate Impact Assessment. 2005; 1:243–352.

Charlesworth D, Charlesworth B. Inbreeding Depression and its Evolutionary Consequences. Annual Review of Ecology and Systematics. 1987; 18(1987):237–268. doi: 10.1146/annurev.es.18.110187.001321.

Clark JS. Why trees migrate so fast: confronting theory with dispersal biology and the paleorecord. The American Naturalist. 1998; 152(2):204–224. doi: 10.1086/286162.

code by Richard A Becker OS, version by Ray Brownrigg Enhancements by Thomas P Minka ARWR, Deckmyn A. maps: Draw Geographical Maps; 2018, https://cran.r-project.org/package=maps.

Corlett RT, Westcott DA. Will plant movements keep up with climate change? Trends in Ecology and Evolution Evolution. 2013 aug; 28(8):482–488. doi: 10.1016/j.tree.2013.04.003.

DeAngelis DL, Mooij WM. Individual-Based Modeling of Ecological and Evolutionary Processes. Annual Review of Ecology, Evolution, and Systematics. 2005; 36:147–168. doi: https://doi.org/10.1146/annurev.ecolsys.36.102003.152644.

Dee DP, Uppala SMM, Simmons AJ, Berrisford P, Poli P, Kobayashi S, Andrae U, Balmaseda MA, Balsamo G, Bauer dP, Others, Bauer P, Bechtold P, Beljaars ACM, van de Berg L, Bidlot J, Bormann N, Delsol C, Dragani R, Fuentes M, >et al. The ERA-Interim reanalysis: Configuration and performance of the data assimilation system. Quarterly Journal of the royal meteorological society. 2011 apr; 137(656):553–597. doi: 10.1002/qj.828.

Epp LS, Kruse S, Kath NJ, Stoof-Leichsenring KR, Tiedemann R, Pestryakova LA, Herzschuh U. Temporal and spatial patterns of mitochondrial haplotype and species distributions in Siberian larches inferred from ancient environmental DNAand modeling. Scientific Reports. 2018 dec; 8(1):17436. doi: 10.1038/s41598-018-35550-w.

Guo W, Rees G. Correlation between the dynamics and spatial configuration of the circumarctic latitudinal forest-tundra ecotone. International Journal of Remote Sensing. 2021; 42(4):1250–1274. doi: 10.1080/01431161.2020.1826062.

Hansson A, Dargusch P, Shulmeister J. A review of modern treeline migration, the factors controlling it and the implications for carbon storage. Journal of Mountain Science. 2021; 18(2):291–306. doi: 10.1007/s11629-020-6221-1.

Harris I, Osborn TJ, Jones P, Lister D. Version 4 of the CRU TS monthly high-resolution gridded multivariate climate dataset. Scientific Data. 2020; 7(1):1–18. doi: 10.1038/s41597-020-0453-3.

Harsch MA, Bader MY. Treeline form - a potential key to understanding treeline dynamics. Global Ecology and Biogeography. 2011; 20:582–596. doi: 10.1111/j.1466-8238.2010.00622.x.

Harsch MA, Hulme PE, McGlone MS, Duncan RP. Are treelines advancing? A global meta-analysis of treeline response to climate warming. Ecology Letters. 2009 oct; 12(10):1040–1049. doi: 10.1111/j.1461-0248.2009.01355.x.

Herzschuh U. Legacy of the Last Glacial on the present-day distribution of deciduous versus evergreen boreal forests. Global Ecology and Biogeography. 2020 feb; 29(2):198–206. doi: 10.1111/geb.13018.

Hijmans RJ. raster: Geographic Data Analysis and Modeling; 2020, https://cran.r-project.org/package=raster.

Hijmans RJ, Cameron SE, Parra JL, Jones PG, Jarvis A. Very high resolution interpolated climate surfaces for global land areas. International Journal of Climatology. 2005 dec; 25(15):1965–1978. doi: 10.1002/joc.1276.

Hofgaard A, Harper KA, Golubeva E. The role of the circumarctic forest-tundra ecotone for Arctic biodiversity. Biodiversity. 2012; 13(3-4):174–181. doi: 10.1080/14888386.2012.700560.

Kaplan JO, Bigelow NH, Prentice IC, Harrison SP, Bartlein PJ, Christensen TR, Cramer W, Matveyeva NV, Mcguire AD, Murray DF. Climate change and Arctic ecosystems: and future projections. Journal of Geophysical Research. 2003; 108(D19):8171. doi: 10.1029/2002JD002559.

Kharuk VI, Ranson KJ, Im ST, Oskorbin PA, Dvinskaya ML, Ovchinnikov DV, Plateau A, Maria L. Tree-Line Structure and Dynamics at the Northern Limit of the Larch Forest: Anabar Plateau, Siberia, Russia. Arctic, Antarctic, and Alpine Research. 2013; 45(4):526–537. doi: 10.1657/1938-4246-45.4.526.

Krever V, Stishov M, Onufrenya I. National protected areas of the Russian Federation: Gap analysis and perspective Framework. Moscow: WWF; 2009.

Kruse S, Stuenzi SM, Boike J, Langer M, Gloy J, Herzschuh U. Novel coupled permafrost-forest model revealing the interplay between permafrost, vegetation, and climate across eastern Siberia. Geoscientific Model Development Discussions. 2021; 2021:1–35. https://gmd.copernicus.org/preprints/gmd-2021-304/, doi: 10.5194/gmd-2021-304.

Kruse S, Gerdes A, Kath NJ, Epp LS, Stoof-Leichsenring KR, Pestryakova LA, Herzschuh U. Dispersal distances and migration rates at the arctic treeline in Siberia – a genetic and simulation-based study. Biogeosciences. 2019 mar; 16(6):1211–1224. doi: 10.5194/bg-16-1211-2019.

Kruse S, Herzschuh U, Stünzi S, Vyse S, Zakharov E. Sampling mixed-species boreal forests affected by disturbances and mountain lake mountain lake and alas lake coring in Central Yakutia. In: Kruse S, Bol-shiyanov D, Grigoriev MN, Morgenstern A, Pestryakova L, Tsibizov L, Udke A, editors. Russian-German Cooperation: Expeditions to Siberia in 2018. Berichte zur Polar-und Meeresforschung = Reports on polar and marine research Bremerhaven: Alfred Wegener Institute for Polar and Marine Research; 2019.p. 148–153. doi: 10.2312/BzPM_0734_2019.

Kruse S, Stoof-Leichsenring KR. Keperveem - Past and present vegetation dynamics at the most eastern extension of the Siberian boreal treeline. In: Overduin PP, Blender F, Bolshiyanov DY, Grigoriev MN, Morgenstern A, Meyer H, editors. Russian-German Cooperation: Expeditions to Siberia in 2016, berichte z ed. Alfred-Wegener-Institut, Helmholtz-Zentrum für Polar-und Meeresforschung, Bremerhaven, Germany: Alfred-Wegener-Institut, Helmholtz-Zentrum für Polar-und Meeresforschung, Bremerhaven, Germany; 2016.p. 130–137. doi: 10.2312/BzPM_0709_2017.

Kruse S, Wieczorek M, Jeltsch F, Herzschuh U. Treeline dynamics in Siberia under changing climates as inferred from an individual-based model for Larix. Ecological Modelling. 2016; 338:101–121. doi: 10.1016/j.ecolmodel.2016.08.003.

Liu S, Stoof-Leichsenring KR, Kruse S, Pestryakova LA, Herzschuh U. Holocene Vegetation and Plant Diversity Changes in the North-Eastern Siberian Treeline Region From Pollen and Sedimentary Ancient DNA. Frontiers in Ecology and Evolution. 2020 sep; 8(September):1–17. doi: 10.3389/fevo.2020.560243.

Loarie SR, Duffy PB, Hamilton H, Asner GP, Field CB, Ackerly DD. The velocity of climate change. Nature. 2009; 462(7276):1052–1055. doi: 10.1038/nature08649.

MacDonald GM, Kremenetski KV, Beilman DW. Climate change and the northern Russian treeline zone. Philosophical Transactions of the Royal Society B: Biological Sciences. 2008 jul; 363(1501):2283–2299. doi: 10.1098/rstb.2007.2200.

Mamet SD, Brown CD, Trant AJ, Laroque CP. Shifting global Larix distributions: Northern expansion and southern retraction as species respond to changing climate. Journal of Biogeography. 2019; 46(September 2018):30–44. doi: 10.1111/jbi.13465.

May F, Giladi I, Ristow M, Ziv Y, Jeltsch F. Metacommunity, mainland-island system or island communities? Assessing the regional dynamics of plant communities in a fragmented landscape. Ecography. 2013 jul; 36(7):842–853. doi: 10.1111/j.1600-0587.2012.07793.x.

Meinshausen M, Smith SJ, Calvin K, Daniel JS, Kainuma MLT, Lamarque J, Matsumoto K, Montzka SA, Raper SCB, Riahi K, Thomson A, Velders GJM, van Vuuren DPP. The RCP greenhouse gas concentrations and their extensions from 1765 to 2300. Climatic Change. 2011; 109(1):213–241. doi: 10.1007/s10584-011-0156-z.

Mod HK, Luoto M. Arctic shrubification mediates the impacts of warming climate on changes to tundra vegetation. Environmental Research Letters. 2016 dec; 11(12):124028. doi: 10.1088/1748-9326/11/12/124028.

Mona S, Ray N, Arenas M, Excoffier L. Genetic consequences of habitat fragmentation during a range expansion. Heredity. 2014; 112(3):291–9. doi: 10.1038/hdy.2013.105.

Morelli TL, Barrows CW, Ramirez AR, Cartwright JM, Ackerly DD, Eaves TD, Ebersole JL, Krawchuk MA, Letcher BH, Mahalovich MF, Meigs GW, Michalak JL, Millar CI, Quiñones RM, Stralberg D, Thorne JH. Climate-change refugia: biodiversity in the slow lane. Frontiers in Ecology and the Environment. 2020 jun; 18(5):228–234. doi: 10.1002/fee.2189.

Ohsawa T, Ide Y. Global patterns of genetic variation in plant species along vertical and horizontal gradients on mountains. Global Ecology and Biogeography. 2008 mar; 17(2):152–163. doi: 10.1111/j.1466-8238.2007.00357.x.

Opel T, Fritzsche D, Meyer H. Eurasian Arctic climate over the past millennium as recorded in theAkademii Nauk ice core (Severnaya Zemlya). Climate of the Past. 2013 oct; 9(5):2379–2389. doi: 10.5194/cp-9-2379-2013.

Pauli H, Gottfried M, Dullinger S, Abdaladze O, Akhalkatsi M, Alonso JLB, Coldea G, Dick J, Erschbamer B, Calzado RF, Ghosn D, Holten JI, Kanka R, Kazakis G, Kollar J, Larsson P, Moiseev P, Moiseev D, Molau U, Mesa JM, et al. Recent Plant Diversity Changes on Europe’s Mountain Summits. Science. 2012 apr; 336(6079):353–355. doi: 10.1126/science.1219033.

Pearson RG, Phillips SJ, Loranty MM, Beck PSAA, Damoulas T, Knight SJ, Goetz SJ. Shifts in Arctic vegetation and associated feedbacks under climate change. Nature Climate Change. 2013 mar; 3(7):673–677. doi: 10.1038/nclimate1858.

Pebesma E. Simple Features for R: Standardized Support for Spatial Vector Data. The RJournal. 2018; 10(1):439–446. doi: 10.32614/RJ-2018-009.

QGIS Development Team. QGIS Geographic Information System. QGIS Association; 2020, https://www.qgis.org.

R Core Team. R: A Language and Environment for Statistical Computing. R Foundation for Statistical Computing, Vienna, Austria; 2019, https://www.r-project.org/.

Rees WG, Hofgaard A, Boudreau S, Cairns DM, Harper K, Mamet S, Mathisen I, Swirad Z, Tutubalina O. Is subarctic forest advance able to keep pace with climate change? Global Change Biology. 2020; 26(7):3965–3977. doi: 10.1111/gcb.15113.

Scherrer D, Vitasse Y, Guisan A, Wohlgemuth T, Lischke H. Competition and demography rather than dispersal limitation slow down upward shifts of trees’ upper elevation limits in the Alps. Journal of Ecology. 2020 nov; 108(6):2416–2430. doi: 10.1111/1365-2745.13451.

Secretariat of the United Nations Convention on Biological Diversity. First Draft of the Post-2020 Global Biodiversity Framework. Cbd/Wg2020/3/3. 2021; (July):1–12. https://www.cbd.int/doc/c/abb5/591f/2e46096d3f0330b08ce87a45/wg2020-03-03-en.pdf.

Shevtsova I, Heim B, Kruse S, Schröder J, Troeva EI, Pestryakova LA, Zakharov ES, Herzschuh U. Strong shrub expansion in tundra-taiga, tree infilling in taiga and stable tundra in central Chukotka (north-eastern Siberia) between 2000 and 2017. Environmental Research Letters. 2020 aug; 15(8):085006. doi: 10.1088/1748-9326/ab9059.

Shevtsova I, Herzschuh U, Heim B, Pestryakova LA, Zakharov ES, Kruse S. Future spatially explicit tree above-ground biomass trajectories revealed for a mountainous treeline ecotone using the individual-based model LAVESI. Environmental Research Letters. 2021; submitted.

Snell RS, Cowling SA. Consideration of dispersal processes and northern refugia can improve our understanding of past plant migration rates in North America. Journal of Biogeography. 2015 sep; 42(9):1677–1688. doi: 10.1111/jbi.12544.

Stralberg D, Arseneault D, Baltzer JL, Barber QE, Bayne EM, Boulanger Y, Brown CD, Cooke HA, Devito K, Edwards J, Estevo CA, Flynn N, Frelich LE, Hogg EH, Johnston M, Logan T, Matsuoka SM, Moore P, Morelli TL, Morissette JL, >et al. Climate-change refugia in boreal North America: what, where, and for how long? Frontiers in Ecology and the Environment. 2020 jun; 18(5):261–270. doi: 10.1002/fee.2188.

Stuenzi SM, Boike J, Cable W, Herzschuh U, Kruse S, Pestryakova LA, Schneider von Deimling T, Westermann S, Zakharov ES, Langer M. Variability of the surface energy balance in permafrost-underlain boreal forest. Biogeosciences. 2021 jan; 18(2):343–365. doi: 10.5194/bg-18-343-2021.

Sullivan PF, Ellison SBZ, McNown RW, Brownlee AH, Sveinbjörnsson B. Evidence of soil nutrient availability as the proximate constraint on growth of treeline trees in northwest Alaska. Ecology. 2015; 96(3):716–727. doi: 10.1890/14-0626.1.

Väliranta M, Kaakinen A, Kuhry P, Kultti S, Salonen JS, Seppä H. Scattered late-glacial and early Holocene tree populations as dispersal nuclei for forest development in north-eastern European Russia. Journal of Biogeography. 2011 may; 38(5):922–932. doi: 10.1111/j.1365-2699.2010.02448.x.

Walker DA, Raynolds MK, Daniëls FJAA, Einarsson E, Elvebakk A, Gould WA, Katenin AE, Kholod SS, Markon CJ, Melnikov ES, Moskalenko NG, Talbot SS, Yurtsev BA, Team tomotC. The circumpolar Arctic vegetation map. Journal of Vegetation Science. 2005; 16(3):267–282. doi: 10.1658/1100-9233(2005)016[0267:TCAVM]2.0.CO;2.

Wieczorek M, Kolmogorov A, Kruse S, Jacobsen I, Nitze I, Nikolaev A, Heinrich I, Pestryakova L, Herzschuh U. Disturbance-effects on treeline larch-stands in the lower Kolyma River area (NE Siberia). Silva Fennica. 2017; 51(3):1–20. doi: 10.14214/sf.1666.

Wieczorek M, Kruse S, Epp LS, Kolmogorov A, Nikolaev AN, Heinrich I, Jeltsch F, Pestryakova LA, Zibulski R, Herzschuh U. Dissimilar responses of larch stands in northern Siberia to increasing temperatures-a field and simulation based study. Ecology. 2017 sep; 98(9):2343–2355. doi: 10.1002/ecy.1887.

Wieczorek M, Kruse S, Epp LS, Kolmogorov A, Nikolaev AN, Heinrich I, Jeltsch F, Pestryakova LA, Zibulski R, Herzschuh U, Field data for larches growing in the Taimyr treeline ecotone. PANGAEA; 2017. doi: 10.1594/PANGAEA.874612.

Wielgolaski FE, Hofgaard A, Holtmeier FK. Sensitivity to environmental change of the treeline ecotone and its associated biodiversity in European mountains. Climate Research. 2017; 73(1-2):151–166. doi: 10.3354/cr01474.

Yu Q, Epstein H, Walker D. Simulating the effects of soil organic nitrogen and grazing on arctic tundra vegetation dynamics on the Yamal Peninsula, Russia. Environmental Research Letters. 2009 oct; 4(4):045027. doi: 10.1088/1748-9326/4/4/045027.

Zurell D, König C, Malchow Ak, Kapitza S, Bocedi G, Travis J, Fandos G. Spatially explicit models for decision-making in animal conservation and restoration. Ecography. 2021; (44):1–16. doi: 10.1111/ecog.05787.

